# Bat cellular immunity varies by year and dietary habit in an increasingly fragmented landscape

**DOI:** 10.1101/2023.05.22.541709

**Authors:** Isabella K. DeAnglis, Benjamin R. Andrews, Lauren R. Lock, Kristin E. Dyer, Dmitriy V. Volokhov, M. Brock Fenton, Nancy B. Simmons, Cynthia J. Downs, Daniel J. Becker

## Abstract

Monitoring the health of wildlife populations is essential in the face of increased agricultural expansion and forest fragmentation. Loss of habitat and habitat degradation can negatively affect an animal’s physiological state, possibly resulting in immunosuppression and increased morbidity or mortality. We sought to determine how fragmentation may differentially impact cellular immunity and infection risk in Neotropical bats species regularly infected with bloodborne pathogens, and to evaluate how effects may vary over time and by dietary habit. We studied common vampire bats (*Desmodus rotundus*), northern yellow-shouldered bats (*Sturnira parvidens*), and Mesoamerican mustached bats (*Pteronotus mesoamericanus*), representing the dietary habits of sanguinivory, frugivory, and insectivory respectively, in northern Belize. We compared estimated total white blood cell counts, leukocyte differentials, and infection status with two blood-borne bacterial pathogens (*Bartonella* spp. and hemoplasmas) of 118 bats captured in a broadleaf, secondary forest over a three-year period (2017-2019) of increasing habitat fragmentation. We found evidence for bat species-specific responses of cellular immunity between years, with neutrophil counts increasing in *D. rotundus*, but decreasing in *S. parvidens* and *P. mesoamericanus* from 2018 to 2019. However, the odds of infection with *Bartonella* spp. and hemoplasma spp. between 2017 and 2019 did not differ between bat species, contrary to our prediction that pathogen prevalence may increase with increased fragmentation. We conclude that each bat species invested differently in cellular immunity in ways that changed over years of increasing fragmentation. We recommend further research on the interactions between habitat fragmentation, cellular immunity, and infection across dietary habits of Neotropical bats for informed management and conservation.

## Introduction

Agriculture expansion fragments and degrades landscapes, reducing suitable habitat available to many terrestrial mammals (Laurance *et al*. 2014; Crooks *et al*. 2017). Tropical areas worldwide are experiencing rapid human population growth and associated agriculture expansion, which is particularly concerning for conservation since the tropics support about two thirds of global biodiversity (Antonelli 2022; Bradshaw *et al*. 2009). Impacts of habitat fragmentation on animal physiology, and ultimately on fitness, depend on phenotypic plasticity and how fragmentation affects specific aspects of an animal’s ecology. Generalists may be less likely to be affected by forest fragmentation than specialists (Ramiadantsoa *et al*. 2018), and some species may be epigenetically primed to take advantage of changing landscapes (Kilvitis *et al*. 2017). Species with specialized niches, including highly specific shelter or dietary needs, are more likely to be affected by the physiological challenges associated with habitat fragmentation, resulting in population declines in such species (Kosydar *et al*. 2014). Conversely, some species may be less affected by or even benefit from fragmentation because of increased access to human-provisioned food resources in the form of crops or livestock (Oro *et al*. 2013). The dietary habit of a species can thus impact how habitat changes associated with fragmentation affect individual physiology (Hinam & Clair 2008). Reduced physiological condition, as a result of habitat fragmentation, can result in immunodeficiency and increased morbidity (Villafuerte *et al*. 1997; Johnstone *et al*. 2012; Seltmann *et al*. 2017). Understanding how fragmentation differentially affects the ability of species to mount immune defenses against pathogens can inform conservation and land management decisions, and contribute significantly to our understanding of pathogen dynamics in complex host communities.

Habitat fragmentation may cause immunosuppression and immunomodulation, resulting in increased susceptibility to infections in wildlife (Aguirre & Tabor 2008). When an individual is experiencing more frequent or intense stressors (Davis *et al*. 2008) or is parasitized (Hernandez *et al*. 2018), plasma glucocorticoid levels can increase, often impairing the host immune response (Coutinho & Chapman 2011; Sapolsky *et al*. 2000). Comparing estimated total white blood cell counts (TWBC) and differential white blood cell counts (DWBC) in mammals under different environmental conditions (i.e.,in areas with differential levels of habitat disturbance and fragmentation) is a low-cost and tractable means for determining how environmental changes impact individual investment in cellular immune defenses (Becker *et al*. 2018b; Schneeberger *et al*. 2013). For example, increased glucocorticoid levels are associated with increased TWBC, neutrophilia, lymphopenia, and eosinopenia across vertebrates (Jain 1986; Davis *et al*. 2008). Further, immunocompromised individuals often experience increased pathogen susceptibility, which can increase TWBC counts and cause neutrophilia, lymphopenia, and monocytosis in vertebrates (Jain 1986; Davis *et al*. 2008; Davis *et al*. 2004). Thus, documenting changes in the leukocyte profile and infection status of wildlife over time can help evaluate how habitat fragmentation impacts host cellular immunity and pathogen spread.

The Neotropics contain the greatest diversity of bats globally, with species that span almost all possible mammal dietary habits (Mickleburgh *et al*. 2002, Fenton 1992). Neotropical bats are important members of tropical forest ecosystems because of their role in seed dispersal, insect predation, and pollination (Kunz *et al*. 2011). Bats are also notable for their potential to spread virulent pathogens to humans, livestock, and other species (Chan *et al*. 2013; Allocati *et al*. 2016). Humans that live in or near fragmented forests in the tropics have greater exposure to bat-borne pathogens, due to their proximity to bat hosts at ecotone roost sites (Rulli *et al*. 2017). Because of their great ecological diversity, Neotropical bats are a valuable system for studying effects of habitat fragmentation on diverse ecological interactions.

In this study, we used Neotropical bats as a model system to ask how forest fragmentation affects cellular immunity and infection status of diverse host species, with the goal of testing how dietary habit impacts the immune response to habitat fragmentation. Specifically, we studied common vampire bats (*Desmodus rotundus*), northern yellow-shouldered bats (*Sturnira parvidens*), and Mesoamerican mustached bats (*Pteronotus mesoamericanus*). These species represent the dietary habits of sanguivores, frugivores, and insectivores respectively (Bobrowiec *et al*. 2015, Ingala *et al*. 2021). All three bats are broadly distributed in Central America and are routinely found in fragmented landscapes (Brändel *et al*. 2020; Alpízar *et al*. 2019; Kraker-Castañeda *et al*. 2016; Herrera et al., 2018); however, their response to fragmentation likely differs owing to their different dietary habits. Across its range, *Desmodus rotundus* capitalizes on domestic animal prey (especially cattle) in fragmented landscapes, and this human-provisioned food source is preferentially selected over wildlife prey (Voigt & Kelm 2006; Ingala *et al*. 2019; Bobrowiec *et al*. 2015). Therefore, we predict that the cellular immunity of *D. rotundus* would not be negatively impacted by forest fragmentation (or even could benefit from livestock prey; Becker *et al*. 2018b). *Sturnira parvidens* is a frugivore that specializes in fruit from early successional plants, which are most abundant in fragmented and disturbed forests (Kraker-Castañeda *et al*. 2016; García-Morales *et al*. 2012; Galindo-González *et al*. 2000). Prior work has found that *S. parvidens* is one of the most abundant bat species in at least some fragmented forests (Herrera *et al*. 2018; Ramírez-Lucho *et al*. 2017). In contrast, *Pteronotus mesoamericanus* may be more vulnerable to increasing habitat fragmentation, as these bats forage for insects using echolocation in dense vegetation within forest interiors (Alpízar *et al*. 2019; Núñez *et al*. 2019). Fragmentation could thus reduce access to forest interiors for bat foraging (Núñez *et al*. 2019), which could function as a stressor that impairs *P. mesoamericanus* immunity.

The likelihood of a bat being exposed to pathogens, and the mode of pathogen exposure, can also depend on the ecological niche of a species (Schneeberger *et al*. 2013). In this study, we focused on how fragmentation affects the prevalence of *Bartonella* spp. and hemotropic *Mycoplasma* spp. (i.e., hemoplasmas), which are common bacterial pathogens in Neotropical bats (Becker *et al*. 2018a; Becker *et al*. 2018b; Becker *et al*. 2020a; Ikeda *et al*. 2017). *Bartonella* spp. are intraerythrocytic and are vectored by hematophagous arthropods, including bat ectoparasites, and may also be transmitted through blood, saliva, or feces to a variety of hosts. In humans, these infections can cause a variety of diseases, including cat-scratch disease, endocarditis, and Carrion’s disease (Jacomo *et al*. 2002). Hemoplasmas are obligate parasites of erythrocytes and are thought to be transmitted by direct contact and possibly also vectored by hematophagous arthropods, including bat ectoparasites (Messick 2004; Cohen *et al*. 2018). Hemoplasmas can cause hemolytic anemia, arthritis, pneumonia, conjunctivitis, infertility and other acute to chronic diseases in humans and other mammals (Messick 2004; Millán *et al*. 2021; Descloux *et al*. 2021). Due to the potential of these pathogens to cause zoonotic infections, which can be life-threatening in humans, understanding how the likelihoods of infection with *Bartonella* spp. and hemoplasmas in bats are affected by habitat fragmentation is important to forecast or prevent future pathogen spillover. Here, we compared the leukocyte profiles and infection states of *D. rotundus, S. parvidens,* and *P. mesoamericanus* inhabiting northern Belize over a three-year period of increasing forest fragmentation. We hypothesized that increased fragmentation would impact the species’ cellular immunity and infection status differentially, according to their dietary habit, with *P. mesoamericanus* experiencing the greatest changes in cellular immunity (i.e., increased TWBC counts, monocytosis, neutrophilia, and lymphopenia) and infection status (i.e., increased odds of infection) and *D. rotundus* experiencing only minor changes in cellular immunity and infection status, due to their foraging ecology. Evaluating the immunological impacts of habitat fragmentation on these bat species is important not only to inform bat conservation efforts, but also to predict how future habitat loss can influence pathogen spread in bats and potentially to humans.

## Methods

### Bat Capture and Sampling

As part of broader ecological, immunological, and epidemiological studies of bats in Belize (Herrera *et al*., 2018; Becker *et al*. 2020a; Becker *et al*. 2020, Becker *et al*. 2021a; Becker *et al*. 2022), we sampled *D. rotundus*, *S. parvidens,* and *P. mesoamericanus* during April to May 2017– 2019 within the Lamanai Archaeological Reserve (LAR) of Orange Walk District, Belize (N 17.76343, W 88.65292). The LAR is a broadleaf, secondary protected forest near the New River Lagoon, for which the surrounding matrix is experiencing increasing fragmentation as land is converted to cropland and cattle pastures (Herrera *et al*. 2018; Ingala *et al*. 2019, Figure 1). As described previously, we used mist nets and harp traps to capture bats along flight paths and occasionally at the exits of roosts from 7 PM until 10 PM (Becker *et a*l. 2020a, Becker *et al*. 2021b). Bats were kept in clean cloth bags prior to processing and were identified and sexed based on morphology (e.g., Reid 1997). Between 3-30 µl of blood was sampled, based on bat body mass, by lancing the propatagial vein with a sterile needle (23–30G) and collected in a heparinized capillary tube. Thin blood smears were prepared on glass slides and stained with Wright–Geimsa (Astral Diagnostics 5316 Quick III Set), and remaining blood was stored on Whatman FTA cards at room temperature. All bats for this study were released following sampling. Field procedures were performed according to guidelines for the safe and humane handling of bats published by of the American Society of Mammalogists (Sikes *et al*. 2016) and were approved by the Institutional Animal Care and Use Committees of the University of Georgia (A2014 04-016-Y3-A5) and American Museum of Natural History (AMNHIACUC-20170403, AMNHIACUC-20180123, AMNHIACUC-20190129). Fieldwork and sampling were authorized by the Belize Forest Department under permits WL/2/1/17(16), WL/2/1/17(19), WL/2/1/18(16), and FD/WL/1/19(09).

**Figure 1.**
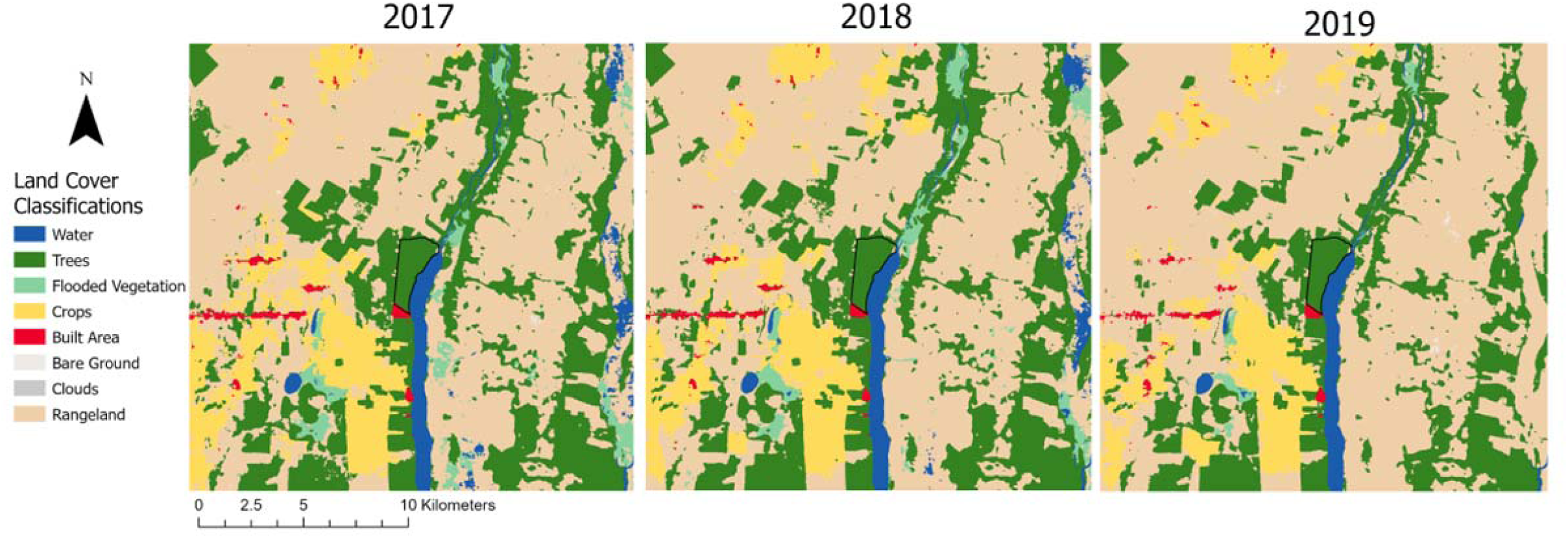
Map of land cover change within 10 km of LAR from 2017 to 2019. Study site in the Orange Walk District in northern Belize, displaying the boundaries of the 450-ha forest of the Lamanai Archaeological Reserve in black (Becker *et al*. 2020). Map and land use/land cover data were derived from the Sentinel-2 10m land use/land cover time series of the world produced by Impact Observatory, Microsoft, and Esri (Karra, Kontgis, et al. 2021). This dataset is based on the dataset produced for the Dynamic World Project by National Geographic Society in partnership with Google and the World Resources Institute.

### White Blood Cell Counts and Statistical Analyses

We estimated TWBC counts for each blood smear (*n* = 118) by averaging the number of leukocytes (neutrophils, lymphocytes, monocytes, basophils, eosinophils) under 10 random fields under 400X magnification (Schneeberger *et al*. 2013). DWBC counts were then estimated by identifying 100 leukocytes and recording the relative abundance of each type of white blood cell under 1000X (oil immersion) magnification. The absolute number of each white blood cell type was then determined by multiplying its relative abundance by the estimated TWBC count (Becker *et al*. 2021a). A subset of the hematology data from 2017 and 2018 were published previously (Becker *et al*. 2021a; Cornelius Ruhs *et al*. 2021).

We used generalized linear models (GLMs) to test how each of our four cellular immunity measures (TWBC, absolute neutrophils, absolute lymphocytes, absolute monocytes) varied among our three species across the three years. We fit separate GLMs with immunity predicted by year, bat species, and their interaction (see Becker *et al*. 2020a for sex effects and other individual-level covariates), modeling each response with a Tweedie distribution (Dunn & Smyth 2005). We used the *mgcv* package in R to fit Tweedie-distributed GLMs using maximum likelihood (Wood 2006). We adjusted for the inflated false-discovery rate in *post-hoc* comparisons with the *emmeans* package (Benjamini & Hochberg 1995).

### Bartonella species and Hemoplasma Infection Analyses

We expanded prior surveys of *Bartonella* spp. and hemoplasmas in Belize with analyses of paired blood samples from 2019 bats with blood smears (Becker *et al*. 2020a; Becker *et al*. 2021a; Becker *et al*. 2018a; Volokhov *et al*. 2017). We extracted DNA from Whatman FTA cards using Qiagen QIAamp DNA Investigator Kits (Volokhov *et al*. 2017). We then used PCR and gel electrophoresis to determine the presence of *Bartonella* spp. (targeting the *gltA* gene) and hemoplasmas (targeting the 16S rRNA gene) with primers and procedures described previously (Volokhov *et al*. 2017; Becker *et al*. 2018a; Becker *et al*. 2020a). We also performed Sanger sequencing analyses for hemoplasmas to assess similarity to our previously established genotypes in Belize bats (see Supplemental Information; Becker *et al*. 2020a).

Following our analyses of WBC data, we derived infection prevalence and 95% confidence intervals (Wilson interval) for each pathogen (and for co-infection) using the *prevalence* package. We then fit GLMs with a binomial distribution per pathogen, with infection status predicted by year, bat species, and their interaction (see Becker *et al*. 2020a for effects of sex and other individual-level covariates on a larger sample size of Belize bat infection status). Because of low sample sizes for pathogen analyses in 2018, we limited this comparison to 2017 and 2019 (*n* = 89). To account for overall smaller sample sizes here, we used the *brglm* package to implement Firth’s bias reduction (Firth 1993).

## Results

### Cellular Immunity

We estimated TWBC and DWBC counts from 42 *D. rotundus*, 40 *S. parvidens,* and 36 *P. mesoamericanus*, in the LAR between 2017 and 2019. Our GLMs found generally strong support for species-specific responses of cellular immunity to year (species-by-year interaction: *F_4_* = 2.04–3.91, *p* = 0.01–0.09; Table 1 and Figure 2). For total leukocytes, the predicted means suggested TWBC of *S. parvidens* increased between 2017 and 2018 and decreased from 2018 to 2019 while TWBC of other species did not change over time, but these *S. parvidens* contrasts were not significant after adjusting for multiple comparisons (Table S1). In contrast, although neutrophil counts did not differ between species in 2017, temporal patterns varied among bats in subsequent years. *D. rotundus* had more counts with increasing fragmentation, whereas neutrophil counts of *S. parvidens* and *P. mesoamericanus* both declined from 2018 to 2019 (Table S2). Lymphocyte counts showed the weakest interactive effect of bat species and year (*F_4_* = 2.04, *p* = 0.09), with *S. parvidens* and *P. mesoamericanus* generally having 1.8 and 2.3 times as many lymphocytes as *D. rotundus* (Table S3). For monocyte counts, predicted means suggested temporal variability in *D. rotundus* and little change for *S. parvidens* or *P. mesoamericanus*, but these contrasts were likewise not significant after adjusting for multiple comparisons (Table S4)

**Figure 2.**
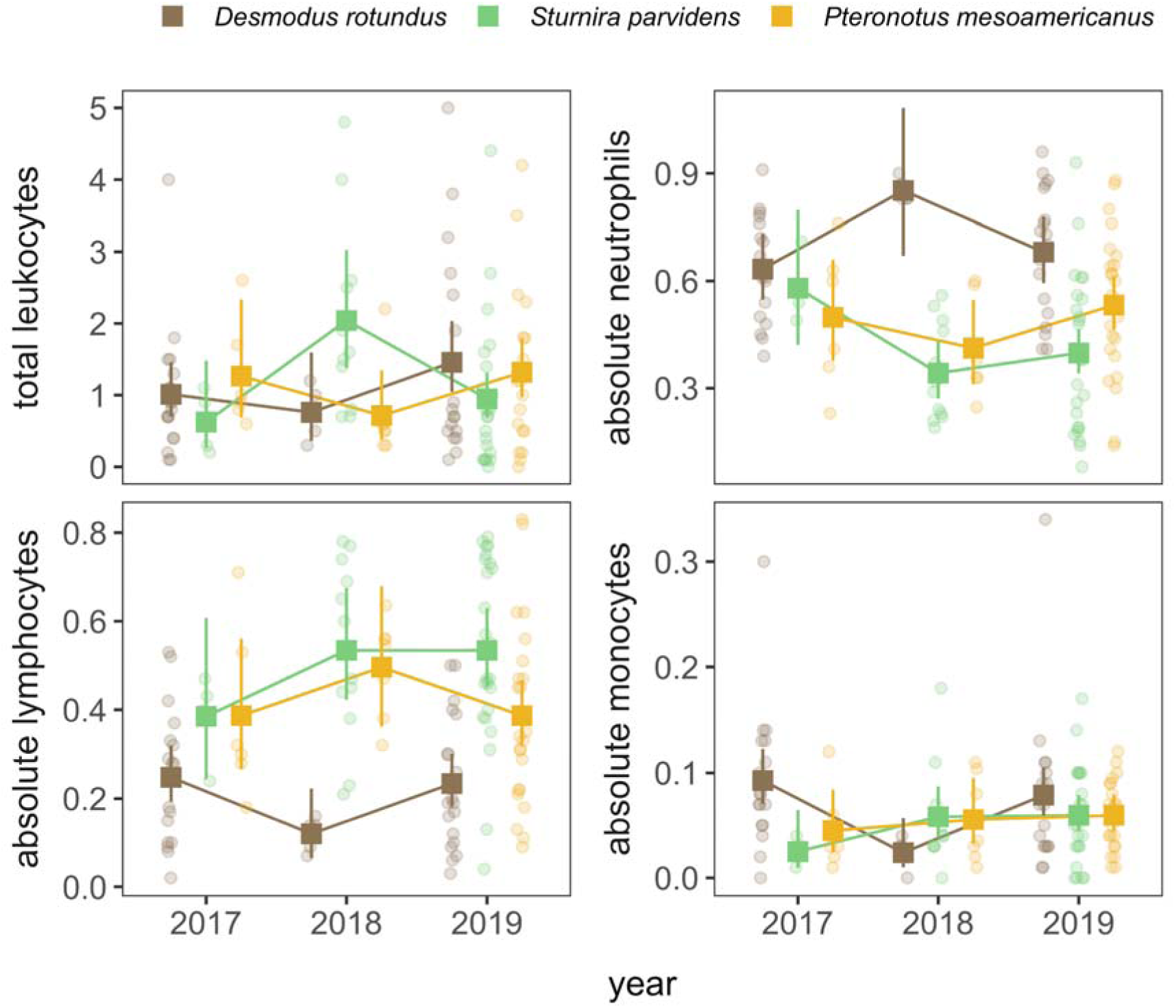
Predicted means and 95% confidence intervals from Tweedie GLMs with interactive effects of bat species and year, for each cellular immunity measure. Raw data are overlaid and jittered.

**Table 1.** Results of Tweedie-distributed GLMs predicting each cellular immunity measure by bat species, year, and their interaction.

| Response | Predictor | *F* | *p* |
| --- | --- | --- | --- |
| Total leukocytes<br>$R^2 = 0.05$ | species | 0.86 | 0.43 |
|  | year | 1.78 | 0.17 |
|  | species*year | 3.91 | <0.01 |
| Absolute neutrophils<br>$R^2 = 0.33$ | species | 1.13 | 0.33 |
|  | year | 2.15 | 0.12 |
|  | species*year | 3.34 | 0.01 |
| Absolute lymphocytes<br>$R^2 = 0.32$ | species | 2.55 | 0.08 |
|  | year | 2.31 | 0.10 |
|  | species*year | 2.04 | 0.09 |
| Absolute monocytes<br>$R^2 = 0.05$ | species | 4.95 | <0.01 |
|  | year | 4.26 | 0.02 |
|  | species*year | 2.78 | 0.03 |

### Blood-borne Pathogen Infections

For the 89 bats screened for bacterial infection in 2017 and 2019 (37 *D. rotundus*, 28 *S. parvidens*, 29 *P. mesoamericanus*), 69% were positive for *Bartonella* spp. (CI: 58–77%), 64% were positive for hemoplasmas (CI: 54–73%), and 45% had coinfections (CI: 35–55%). Our GLMs revealed no effects of species, year, or their interaction on the probability of infection for either *Bartonella* spp. or hemoplasmas (Table 2). Such results thereby suggest little shift in these potentially chronic infections over time on a per-species basis (Figure 3), although such conclusions may be limited by the smaller sample sizes here.

**Figure 3.**
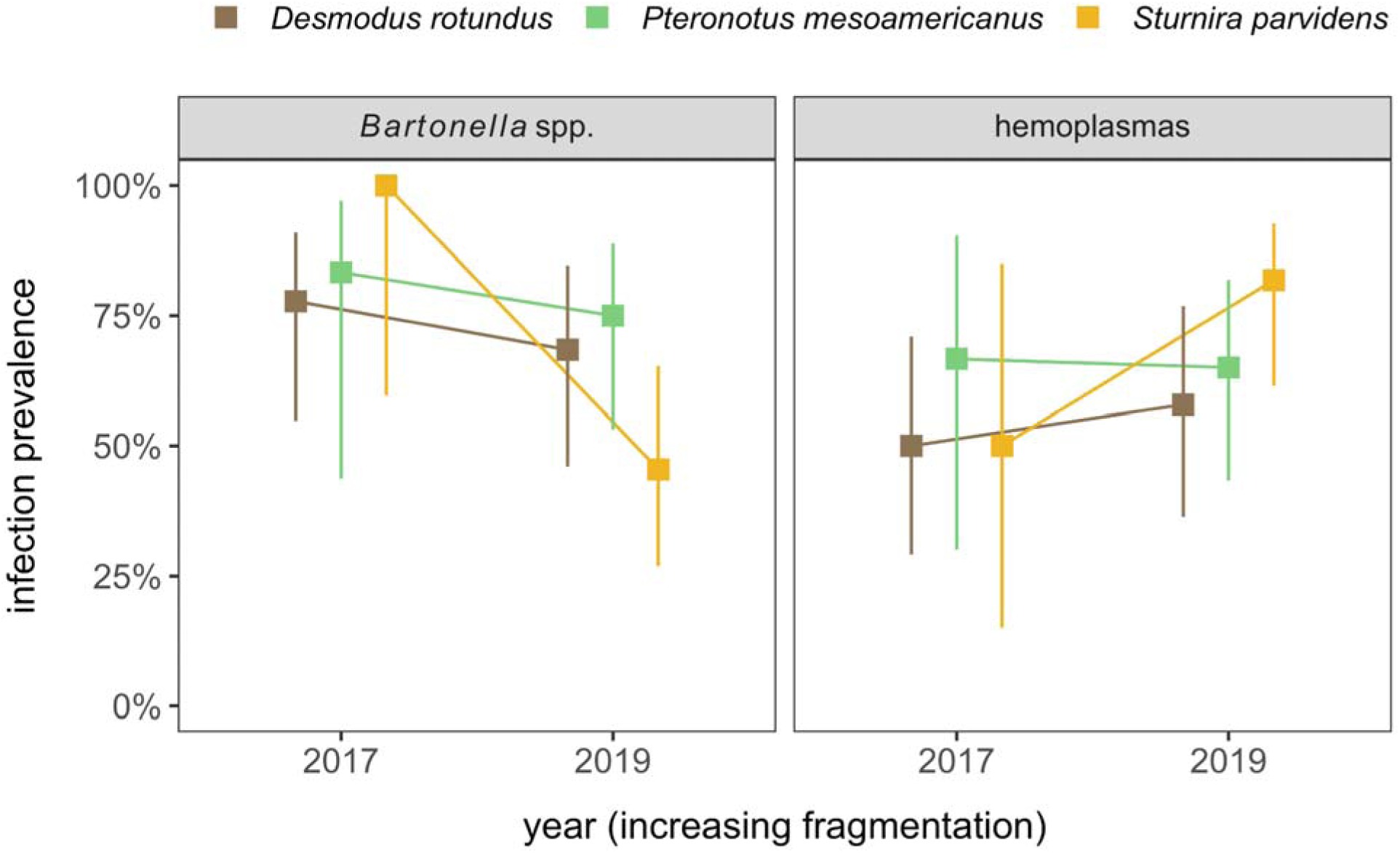
Prevalence of *Bartonella* spp. and hemoplasma infection alongside 95% confidence intervals (Wilson’s interval) stratified by bat species and year.

**Table 2.** Results of binomial GLMs predicting *Bartonella* spp. or hemoplasma infection status by bat species, year, and their interaction. Reference levels are *D. rotundus* for bat species and 2017 for year.

| Response | Coefficient | OR | $z$ | $p$ |
| --- | --- | --- | --- | --- |
| *Bartonella*<br>spp.<br>$R^2 = 0.09$ | intercept | 3.22 | 2.11 | 0.03 |
|  | *P. mesoamericanus* | 1.14 | 0.11 | 0.91 |
|  | *S. parvidens* | 2.79 | 0.58 | 0.56 |
|  | 2019 | 0.64 | -0.59 | 0.55 |
|  | *P. mesoamericanus* : 2019 | 1.19 | 0.13 | 0.90 |
|  | *S. parvidens* : 2019 | 0.14 | -1.03 | 0.30 |
| Hemoplasmas<br>$R^2 = 0.06$ | intercept | 1.00 | 0.00 | 1.00 |
|  | *P. mesoamericanus* | 1.80 | 0.60 | 0.55 |
|  | *S. parvidens* | 1.00 | 0.00 | 1.00 |
|  | 2019 | 1.35 | 0.46 | 0.65 |
|  | *P. mesoamericanus* : 2019 | 0.74 | -0.26 | 0.80 |
|  | *S. parvidens* : 2019 | 3.03 | 0.85 | 0.40 |

Sequencing of hemoplasma positives from 2019 updated a long-term dataset of the diversity of these pathogens in our study area (Volokhov *et al*. 2017; Becker *et al*. 2020a; see Supplemental Materials for more information). We confirmed previously established genotypes in each of our three species (*D. rotundus*: VBG1, VBG2, VBG3; *S. parvidens*: SP1 groups A–C; *P. mesoamericanus*: PPM1). One *S. parvidens* had a novel genotype (SP2, GenBank accession OQ308927) showing 97% similarity to the APH3 genotype, observed only in *Artibeus intermedius* (Becker *et al*. 2020a). *P. mesoamericanus* was also infected with the SP1 genotype and with new *Pteronotus*-specific genotype (PPM2, GenBank accession OQ308895; 97% similar to VBG1). Lastly, *P. mesoamericanus* was also infected by a novel non-hemotropic *Mycoplasma*, the *M. moatsii*–like genotype 4 (GenBank accession OQ308889). All 16S rRNA sequence data are available on GenBank through accession OQ308885–7, OQ308889, OQ308893, OQ308895–97, OQ308899–900, OQ308902–12, OQ308914–19, OQ308923–24, OQ308926–27, OQ533048, OQ546518–19, OQ546527–28, OQ546533, OQ546547, OQ546552, and OQ546555.

## Discussion

By examining changes in leukocyte profiles and bloodborne pathogen (*Bartonella* spp. and hemoplasmas) prevalence in bat species representing distinct dietary habits over years of increasing habitat fragmentation, we expanded our understanding of how anthropogenic habitat modification may differentially affect immunity and infection prevalence of bats. During a period of increasing habitat fragmentation between 2017 and 2019, we found that *D. rotundus, S. parvidens*, and *P. mesoamericanus* exhibited differences in cellular immune defenses. These findings support our prediction that hosts belonging to distinct dietary habits are differentially affected by a changing landscape over time and that these impacts manifest in changes to immune strategy. However, within this sample of bats, we found that *Bartonella* spp. and hemoplasma infection risk did not deviate among years or species. Interestingly, temporal and dietary differences in bat cellular immunity did not translate into variation in *Bartonella* spp. nor hemoplasma risk, at least not over the time scale of this study.

Hosts invest differently in their immune systems based on the varied costs associated with cellular and adaptive immunity (Chaplin 2010). Although investing more energy into adaptive immunity may be more energetically costly during development, cellular immunity can also be costly to maintain later in life when an individual acquires a pathogen (Klasing 2004; McDade *et al*. 2016). Among other factors, life history traits (Lochmiller & Deerenberg 2000), lifespan (Previtali *et al*. 2012) and resource availability (Becker *et al*. 2018b) can influence immune investment. Thus, it is likely that species of bats with different ecological niches will show distinct immune investment strategies. For example, a study of the bat community at our study site previously found species-specific differences in the relationship between mercury and cellular immunity, wherein bats that rely on aquatic prey and bats in agricultural habitats had higher mercury levels and neutropenia as compared to bats of other ecological niches (Becker *et al*. 2021a). Therefore, the distinct ecological requirements of *D. rotundus, S. parvidens, and P. mesoamericanus* may drive species-level variation in how these bats invest in immunity (and in turn how this is affected by habitat fragmentation).

Temporal patterns of immune investment, as measured by leukocyte profiles, varied among bat species, which could at least partially arise from differences in dietary habits. Because diet will likely impact the probability of acquiring a pathogen (Han *et al*. 2021) and can also directly shape immune phenotypes (Schneeberger *et al*. 2013), bat dietary habits will likely affect immune changes over time. Sanguivorous bats are likely to acquire pathogens by ingesting infected blood from other vertebrates, insectivorous bats may acquire pathogens by ingesting infected insects, while frugivorous bats may be more likely to acquire pathogens by ingesting fruit contaminated with feces or saliva (Schneeberger *et al*. 2013). Differences in dietary habits may explain why neutrophil counts of *D. rotundus* increased between 2018 and 2019, while neutrophil counts of *S. parvidens* and *P. mesoamericanus* declined. If *D. rotundus* encounters more pathogens when feeding on infected blood, it may be beneficial for them to increase investment in cellular immunity (i.e., increased neutrophil count) to avoid mounting an energetically costly immune response during periods of increased stress, such as what might be caused by habitat degradation, roost disturbance, or forest fragmentation (Read & Allen 2000; Allen *et al*. 2008).

Other factors that may influence patterns observed in bat leukocyte profiles among species include sociality and roosting behavior (Calisher et al. 2006; Mühldorfer et al. 2011). Both *D. rotundus* and *P. mesoamericanus* are highly social species that live in colonies of a few thousand individuals (Becker *et al*. 2020a; Santana *et al*. 2011; Wilkinson 1986; Wilkinson 1985; Clement & Kanwal 2012), while *S. parvidens* roost alone or in groups of up to 10 individuals (Fenton et al. 2006; Evelyn & Styles 2003). Highly social species such as *D. rotundus* and *P. mesoamericanus* may experience greater pathogen risk compared to less social species such as *S. parvidens* (Stanko *et al*. 2002; Webber & Willis 2016). Moreover, roosts that are permanent and protected from precipitation, such as caves, have increased infection risk compared to more ephemeral or less protected roosts, such as tree cavities (Patterson *et al*. 2007). Therefore, species such as *D. rotundus* and *P. mesoamericanus,* which roost more commonly in caves, could have increased infection risk compared to bats that only inhabit caves rarely and only at night for brief periods (e.g., *S. parvidens*; Sapey 2019). Variation in infection risk as a result of sociality, roosting behavior, and/or other factors is likely to affect individual investment in cellular immunity over time and contribute to the species-specific responses of cellular immunity we observed (Patterson *et al*. 2007; Allen *et al*. 2008; Schneeberger *et al*. 2013).

Our findings suggest different species of bats experience changes in cellular immune investment over time, some of which may be linked to habitat fragmentation. However, we did not find evidence for these changes being associated with increased risk of pathogen infection. Absence of significant changes in infection may be attributed to similar rates of pathogen exposure over time and/or low levels of *Bartonella* spp. and hemoplasmas in the blood. Additionally, the ability to feed on diverse prey (*D. rotundus*) or select early-successional habitats for preferred fruit (*S. parvidens*) may buffer these bats from stressors associated with habitat fragmentation. Although the matrix beyond the LAR is becoming increasingly fragmented (Herrera *et al*. 2018; Ingala *et al*. 2019, Figure. 1), our study site remains a relatively large and protected habitat and as such contains mature trees and caves for roosting. If prey availability, caves, and mature forests persist in the LAR, bats that rely on these dietary and shelter niches may be able to persist without experiencing increased pathogen exposure from habitat reduction.

Although our findings suggest that our select bat species invested differently in immune defenses and that temporal patterns of immune investment varied among species during a period of increasing fragmentation, further research is needed to establish the role of dietary habits in shaping such trends. We selected one representative species per dietary habit for this analysis, but replication is needed to better establish the role of foraging ecology relative to bat taxonomy and other species traits. For example, roosting behavior, body size, sociality, and preferred habitat type are other factors that may influence how fragmentation affects bats and their immunity (Ávila-Gómez et al. 2015, Schneeberger et al. 2013). Comparisons of other frugivorous and insectivorous species, across periods of land conversion would be highly informative, given the rarity of sanguivory among bats. Similarly, our single included insectivore (*P. mesoamericanus*) belongs to the sister family (Mormoopidae) of the other two species (members of Phyllostomidae); inclusion of insectivorous or nectarivorous and/or omnivorous phyllostomids would facilitate more comprehensive comparisons within a clade.

Due to the important ecological roles played by bats in Neotropical ecosystems (e.g., seed dispersal, pollination), understanding how fragmentation impacts bat immunity is critical to conservation of both species and ecosystem function. Frugivorous bats, such as *S. parvidens*, play essential roles in forest regeneration in Neotropical ecosystems (Mello *et al*. 2008), while insectivorous bats, such as *P. mesoamericanus*, are important predators of insects (Alpízar et al. 2019; Núñez et al. 2019), including families that are considered agricultural pests (Ingala *et al*. 2021). Likewise, the preservation of mature trees (the preferred roosting habitat of *S. parvidens*), is essential for the conservation of *S. parvidens* (Evelyn and Stiles 2003; Galindo-González *et al*. 2000). When fragmentation reduces access to mature trees, the physiological condition of these bats could suffer, and, consequently, result in increased stressors and/or reduced immunocompetence. Similarly, *P. mesoamericanus* requires dense forest interiors to forage for insects, which makes them particularly vulnerable to the immunological stressors of major habitat loss and fragmentation (Alpízar et al. 2019; Núñez et al. 2019). In terms of conservation, while *S. parvidens* and *P. mesoamericanus* are of Least Concern status according to the IUCN (Solari 2019; Solari 2016), they are susceptible to extirpation in landscapes void of dense, mature forest. Thus, we suggest monitoring these species in areas experiencing increasing fragmentation (Hernández-Canchola & León-Paniagua, 2020; Evelyn and Stiles 2003; Galindo-González et al. 2000). Lastly, understanding how habitat fragmentation impacts the immunity of *D. rotundus* is important for monitoring rabies virus transmission and control strategies throughout the Neotropics (Becker *et al*. 2020b).

In sum, our study provides evidence that *D. rotundus*, *S. parvidens*, and *P. mesoamericanus* invested differently in cellular immunity in ways that changed over years of increasing fragmentation, suggesting that investment in immune defenses varies by species and dietary habit. Despite this, we found no evidence that habitat fragmentation affected *Bartonella* spp. or hemoplasma infection risk in our increasingly fragmented site, although comparisons were limited by small sample sizes and the short time span of our study. Comparisons of infection prevalence of bats between habitat undergoing rapid fragmentation and habitat remaining intact should be made to determine the role of habitat fragmentation in shaping infection risk. Studying the immunological effects of fragmentation on Neotropical species is important for wildlife management, disease management, and conservation, and continual monitoring of the bat species included in this study is recommended as their habitat becomes increasingly fragmented.

## Acknowledgements

For assistance with field logistics, bat sampling, and permits, we thank Mark Howells, Neil Duncan, and staff of the Lamanai Field Research Center, as well as many colleagues who helped net bats during 2017– 2019 research in Belize. We also thank Eaqan Chaudhry and Konstantin Chumakov for laboratory support.

