## Supplemental Materials for "Bat cellular immunity varies by year and dietary habit in an increasingly fragmented landscape"

Table S1. Contrasts from the TWBC GLM

Table S2. Contrasts from the neutrophil count GLM

Table S3. Contrasts from the lymphocyte count GLM

Table S4. Contrasts from the monocyte count GLM

Additional methods and results for hemoplasma diagnostics
Table S1. Contrasts from the TWBC GLM, adjusted for multiple comparisons (Benjamini–Hochberg)

| **Contrast** | **Ratio** | **SE** | ***t*** | ***p*** |
| --- | --- | --- | --- | --- |
| year2017 D. rotundus / year2018 D. rotundus | 1.33 | 0.56 | 0.68 | 0.67 |
| year2017 D. rotundus / year2019 D. rotundus | 0.69 | 0.18 | -1.44 | 0.39 |
| year2017 D. rotundus / year2017 P. mesoamericanus | 0.8 | 0.29 | -0.62 | 0.69 |
| year2017 D. rotundus / year2018 P. mesoamericanus | 1.42 | 0.53 | 0.93 | 0.56 |
| year2017 D. rotundus / year2019 P. mesoamericanus | 0.77 | 0.19 | -1.08 | 0.51 |
| year2017 D. rotundus / year2017 S. parvidens | 1.62 | 0.77 | 1.01 | 0.52 |
| year2017 D. rotundus / year2018 S. parvidens | 0.5 | 0.14 | -2.55 | 0.14 |
| year2017 D. rotundus / year2019 S. parvidens | 1.06 | 0.27 | 0.25 | 0.85 |
| year2018 D. rotundus / year2019 D. rotundus | 0.52 | 0.22 | -1.57 | 0.36 |
| year2018 D. rotundus / year2017 P. mesoamericanus | 0.6 | 0.29 | -1.04 | 0.51 |
| year2018 D. rotundus / year2018 P. mesoamericanus | 1.06 | 0.53 | 0.12 | 0.91 |
| year2018 D. rotundus / year2019 P. mesoamericanus | 0.58 | 0.24 | -1.35 | 0.39 |
| year2018 D. rotundus / year2017 S. parvidens | 1.22 | 0.71 | 0.34 | 0.83 |
| year2018 D. rotundus / year2018 S. parvidens | 0.37 | 0.16 | -2.31 | 0.17 |
| year2018 D. rotundus / year2019 S. parvidens | 0.8 | 0.33 | -0.54 | 0.73 |
| year2019 D. rotundus / year2017 P. mesoamericanus | 1.15 | 0.41 | 0.4 | 0.81 |
| year2019 D. rotundus / year2018 P. mesoamericanus | 2.04 | 0.75 | 1.95 | 0.32 |
| year2019 D. rotundus / year2019 P. mesoamericanus | 1.1 | 0.26 | 0.43 | 0.8 |
| year2019 D. rotundus / year2017 S. parvidens | 2.33 | 1.1 | 1.79 | 0.33 |
| year2019 D. rotundus / year2018 S. parvidens | 0.71 | 0.19 | -1.28 | 0.39 |
| year2019 D. rotundus / year2019 S. parvidens | 1.53 | 0.36 | 1.8 | 0.33 |
| year2017 P. mesoamericanus / year2018 P. mesoamericanus | 1.77 | 0.8 | 1.27 | 0.39 |
| year2017 P. mesoamericanus / year2019 P. mesoamericanus | 0.96 | 0.34 | -0.12 | 0.91 |
| year2017 P. mesoamericanus / year2017 S. parvidens | 2.03 | 1.09 | 1.31 | 0.39 |
| year2017 P. mesoamericanus / year2018 S. parvidens | 0.62 | 0.23 | -1.29 | 0.39 |
| year2017 P. mesoamericanus / year2019 S. parvidens | 1.33 | 0.47 | 0.81 | 0.6 |
| year2018 P. mesoamericanus / year2019 P. mesoamericanus | 0.54 | 0.2 | -1.7 | 0.33 |
| year2018 P. mesoamericanus / year2017 S. parvidens | 1.14 | 0.62 | 0.24 | 0.85 |
| year2018 P. mesoamericanus / year2018 S. parvidens | 0.35 | 0.13 | -2.76 | 0.12 |
| year2018 P. mesoamericanus / year2019 S. parvidens | 0.75 | 0.27 | -0.78 | 0.6 |
| year2019 P. mesoamericanus / year2017 S. parvidens | 2.11 | 0.99 | 1.6 | 0.36 |
| year2019 P. mesoamericanus / year2018 S. parvidens | 0.65 | 0.17 | -1.71 | 0.33 |
| year2019 P. mesoamericanus / year2019 S. parvidens | 1.39 | 0.32 | 1.44 | 0.39 |
| year2017 S. parvidens / year2018 S. parvidens | 0.31 | 0.15 | -2.45 | 0.14 |
| year2017 S. parvidens / year2019 S. parvidens | 0.66 | 0.31 | -0.89 | 0.56 |
| year2018 S. parvidens / year2019 S. parvidens | 2.15 | 0.56 | 2.94 | 0.12 |

Table S2. Contrasts from the neutrophil GLM, adjusted for multiple comparisons (Benjamini–Hochberg)

| **Contrast** | **Ratio** | **SE** | ***t*** | ***p*** |
| --- | --- | --- | --- | --- |
| year2017 D. rotundus / year2018 D. rotundus | 0.74 | 0.11 | -2.07 | 0.09 |
| year2017 D. rotundus / year2019 D. rotundus | 0.93 | 0.09 | -0.71 | 0.55 |
| year2017 D. rotundus / year2017 P. mesoamericanus | 1.27 | 0.2 | 1.49 | 0.2 |
| year2017 D. rotundus / year2018 P. mesoamericanus | 1.54 | 0.25 | 2.65 | 0.03 |
| year2017 D. rotundus / year2019 P. mesoamericanus | 1.19 | 0.12 | 1.69 | 0.16 |
| year2017 D. rotundus / year2017 S. parvidens | 1.09 | 0.2 | 0.49 | 0.67 |
| year2017 D. rotundus / year2018 S. parvidens | 1.85 | 0.26 | 4.37 | <0.01 |
| year2017 D. rotundus / year2019 S. parvidens | 1.59 | 0.17 | 4.28 | <0.01 |
| year2018 D. rotundus / year2019 D. rotundus | 1.25 | 0.18 | 1.6 | 0.18 |
| year2018 D. rotundus / year2017 P. mesoamericanus | 1.71 | 0.32 | 2.85 | 0.02 |
| year2018 D. rotundus / year2018 P. mesoamericanus | 2.07 | 0.39 | 3.84 | <0.01 |
| year2018 D. rotundus / year2019 P. mesoamericanus | 1.6 | 0.23 | 3.32 | 0.01 |
| year2018 D. rotundus / year2017 S. parvidens | 1.47 | 0.3 | 1.89 | 0.11 |
| year2018 D. rotundus / year2018 S. parvidens | 2.49 | 0.43 | 5.32 | <0.01 |
| year2018 D. rotundus / year2019 S. parvidens | 2.14 | 0.31 | 5.21 | <0.01 |
| year2019 D. rotundus / year2017 P. mesoamericanus | 1.37 | 0.22 | 1.96 | 0.1 |
| year2019 D. rotundus / year2018 P. mesoamericanus | 1.65 | 0.26 | 3.13 | 0.01 |
| year2019 D. rotundus / year2019 P. mesoamericanus | 1.28 | 0.13 | 2.47 | 0.04 |
| year2019 D. rotundus / year2017 S. parvidens | 1.17 | 0.21 | 0.9 | 0.44 |
| year2019 D. rotundus / year2018 S. parvidens | 1.99 | 0.28 | 4.96 | <0.01 |
| year2019 D. rotundus / year2019 S. parvidens | 1.71 | 0.18 | 5.08 | <0.01 |
| year2017 P. mesoamericanus / year2018 P. mesoamericanus | 1.21 | 0.25 | 0.93 | 0.44 |
| year2017 P. mesoamericanus / year2019 P. mesoamericanus | 0.94 | 0.15 | -0.42 | 0.69 |
| year2017 P. mesoamericanus / year2017 S. parvidens | 0.86 | 0.19 | -0.7 | 0.55 |
| year2017 P. mesoamericanus / year2018 S. parvidens | 1.45 | 0.27 | 2.01 | 0.09 |
| year2017 P. mesoamericanus / year2019 S. parvidens | 1.25 | 0.2 | 1.37 | 0.24 |
| year2018 P. mesoamericanus / year2019 P. mesoamericanus | 0.77 | 0.12 | -1.6 | 0.18 |
| year2018 P. mesoamericanus / year2017 S. parvidens | 0.71 | 0.15 | -1.57 | 0.18 |
| year2018 P. mesoamericanus / year2018 S. parvidens | 1.2 | 0.23 | 0.99 | 0.42 |
| year2018 P. mesoamericanus / year2019 S. parvidens | 1.03 | 0.17 | 0.21 | 0.84 |
| year2019 P. mesoamericanus / year2017 S. parvidens | 0.92 | 0.16 | -0.48 | 0.67 |
| year2019 P. mesoamericanus / year2018 S. parvidens | 1.56 | 0.22 | 3.18 | 0.01 |
| year2019 P. mesoamericanus / year2019 S. parvidens | 1.34 | 0.14 | 2.74 | 0.02 |
| year2017 S. parvidens / year2018 S. parvidens | 1.69 | 0.34 | 2.6 | 0.03 |
| year2017 S. parvidens / year2019 S. parvidens | 1.46 | 0.26 | 2.07 | 0.09 |
| year2018 S. parvidens / year2019 S. parvidens | 0.86 | 0.12 | -1.06 | 0.39 |

Table S3. Contrasts from the lymphocyte GLM, adjusted for multiple comparisons (Benjamini–Hochberg)

| **Contrast** | **Ratio** | **SE** | ***t*** | ***p*** |
| --- | --- | --- | --- | --- |
| year2017 D. rotundus / year2018 D. rotundus | 2.07 | 0.71 | 2.13 | 0.07 |
| year2017 D. rotundus / year2019 D. rotundus | 1.06 | 0.19 | 0.32 | 0.84 |
| year2017 D. rotundus / year2017 P. mesoamericanus | 0.64 | 0.15 | -1.93 | 0.1 |
| year2017 D. rotundus / year2018 P. mesoamericanus | 0.5 | 0.1 | -3.36 | <0.01 |
| year2017 D. rotundus / year2019 P. mesoamericanus | 0.64 | 0.1 | -2.75 | 0.02 |
| year2017 D. rotundus / year2017 S. parvidens | 0.65 | 0.17 | -1.65 | 0.17 |
| year2017 D. rotundus / year2018 S. parvidens | 0.46 | 0.08 | -4.37 | <0.01 |
| year2017 D. rotundus / year2019 S. parvidens | 0.47 | 0.07 | -4.98 | <0.01 |
| year2018 D. rotundus / year2019 D. rotundus | 0.51 | 0.17 | -1.96 | 0.1 |
| year2018 D. rotundus / year2017 P. mesoamericanus | 0.31 | 0.11 | -3.18 | 0.01 |
| year2018 D. rotundus / year2018 P. mesoamericanus | 0.24 | 0.09 | -4 | <0.01 |
| year2018 D. rotundus / year2019 P. mesoamericanus | 0.31 | 0.1 | -3.54 | <0.01 |
| year2018 D. rotundus / year2017 S. parvidens | 0.31 | 0.12 | -2.97 | 0.01 |
| year2018 D. rotundus / year2018 S. parvidens | 0.22 | 0.08 | -4.42 | <0.01 |
| year2018 D. rotundus / year2019 S. parvidens | 0.22 | 0.07 | -4.56 | <0.01 |
| year2019 D. rotundus / year2017 P. mesoamericanus | 0.61 | 0.14 | -2.19 | 0.07 |
| year2019 D. rotundus / year2018 P. mesoamericanus | 0.47 | 0.1 | -3.66 | <0.01 |
| year2019 D. rotundus / year2019 P. mesoamericanus | 0.61 | 0.1 | -3.13 | 0.01 |
| year2019 D. rotundus / year2017 S. parvidens | 0.61 | 0.16 | -1.87 | 0.11 |
| year2019 D. rotundus / year2018 S. parvidens | 0.44 | 0.08 | -4.72 | <0.01 |
| year2019 D. rotundus / year2019 S. parvidens | 0.44 | 0.07 | -5.39 | <0.01 |
| year2017 P. mesoamericanus / year2018 P. mesoamericanus | 0.78 | 0.19 | -1 | 0.41 |
| year2017 P. mesoamericanus / year2019 P. mesoamericanus | 1 | 0.21 | 0 | 1 |
| year2017 P. mesoamericanus / year2017 S. parvidens | 1 | 0.3 | 0.01 | 1 |
| year2017 P. mesoamericanus / year2018 S. parvidens | 0.72 | 0.16 | -1.44 | 0.23 |
| year2017 P. mesoamericanus / year2019 S. parvidens | 0.72 | 0.15 | -1.55 | 0.19 |
| year2018 P. mesoamericanus / year2019 P. mesoamericanus | 1.28 | 0.24 | 1.33 | 0.26 |
| year2018 P. mesoamericanus / year2017 S. parvidens | 1.29 | 0.36 | 0.89 | 0.46 |
| year2018 P. mesoamericanus / year2018 S. parvidens | 0.93 | 0.19 | -0.37 | 0.83 |
| year2018 P. mesoamericanus / year2019 S. parvidens | 0.93 | 0.17 | -0.41 | 0.82 |
| year2019 P. mesoamericanus / year2017 S. parvidens | 1 | 0.25 | 0.02 | 1 |
| year2019 P. mesoamericanus / year2018 S. parvidens | 0.72 | 0.11 | -2.1 | 0.08 |
| year2019 P. mesoamericanus / year2019 S. parvidens | 0.72 | 0.09 | -2.51 | 0.03 |
| year2017 S. parvidens / year2018 S. parvidens | 0.72 | 0.19 | -1.25 | 0.28 |
| year2017 S. parvidens / year2019 S. parvidens | 0.72 | 0.18 | -1.32 | 0.26 |
| year2018 S. parvidens / year2019 S. parvidens | 1 | 0.15 | 0 | 1 |

Table S4. Contrasts from the monocyte GLM, adjusted for multiple comparisons (Benjamini–Hochberg)

| **Contrast** | **Ratio** | **SE** | ***t*** | ***p*** |
| --- | --- | --- | --- | --- |
| year2017 D. rotundus / year2018 D. rotundus | 3.87 | 1.79 | 2.91 | 0.14 |
| year2017 D. rotundus / year2019 D. rotundus | 1.18 | 0.24 | 0.8 | 0.57 |
| year2017 D. rotundus / year2017 P. mesoamericanus | 2.06 | 0.72 | 2.07 | 0.21 |
| year2017 D. rotundus / year2018 P. mesoamericanus | 1.67 | 0.51 | 1.66 | 0.24 |
| year2017 D. rotundus / year2019 P. mesoamericanus | 1.56 | 0.32 | 2.17 | 0.19 |
| year2017 D. rotundus / year2017 S. parvidens | 3.71 | 1.88 | 2.59 | 0.14 |
| year2017 D. rotundus / year2018 S. parvidens | 1.59 | 0.4 | 1.86 | 0.23 |
| year2017 D. rotundus / year2019 S. parvidens | 1.56 | 0.31 | 2.2 | 0.19 |
| year2018 D. rotundus / year2019 D. rotundus | 0.3 | 0.14 | -2.56 | 0.14 |
| year2018 D. rotundus / year2017 P. mesoamericanus | 0.53 | 0.29 | -1.15 | 0.41 |
| year2018 D. rotundus / year2018 P. mesoamericanus | 0.43 | 0.22 | -1.62 | 0.24 |
| year2018 D. rotundus / year2019 P. mesoamericanus | 0.4 | 0.19 | -1.95 | 0.21 |
| year2018 D. rotundus / year2017 S. parvidens | 0.96 | 0.63 | -0.06 | 0.98 |
| year2018 D. rotundus / year2018 S. parvidens | 0.41 | 0.2 | -1.82 | 0.23 |
| year2018 D. rotundus / year2019 S. parvidens | 0.4 | 0.19 | -1.96 | 0.21 |
| year2019 D. rotundus / year2017 P. mesoamericanus | 1.75 | 0.62 | 1.6 | 0.24 |
| year2019 D. rotundus / year2018 P. mesoamericanus | 1.42 | 0.44 | 1.13 | 0.41 |
| year2019 D. rotundus / year2019 P. mesoamericanus | 1.32 | 0.27 | 1.35 | 0.32 |
| year2019 D. rotundus / year2017 S. parvidens | 3.16 | 1.6 | 2.26 | 0.19 |
| year2019 D. rotundus / year2018 S. parvidens | 1.35 | 0.34 | 1.2 | 0.4 |
| year2019 D. rotundus / year2019 S. parvidens | 1.32 | 0.27 | 1.37 | 0.32 |
| year2017 P. mesoamericanus / year2018 P. mesoamericanus | 0.81 | 0.34 | -0.5 | 0.77 |
| year2017 P. mesoamericanus / year2019 P. mesoamericanus | 0.75 | 0.27 | -0.8 | 0.57 |
| year2017 P. mesoamericanus / year2017 S. parvidens | 1.8 | 1.05 | 1.01 | 0.47 |
| year2017 P. mesoamericanus / year2018 S. parvidens | 0.77 | 0.29 | -0.68 | 0.64 |
| year2017 P. mesoamericanus / year2019 S. parvidens | 0.76 | 0.26 | -0.8 | 0.57 |
| year2018 P. mesoamericanus / year2019 P. mesoamericanus | 0.93 | 0.29 | -0.23 | 0.96 |
| year2018 P. mesoamericanus / year2017 S. parvidens | 2.22 | 1.24 | 1.43 | 0.31 |
| year2018 P. mesoamericanus / year2018 S. parvidens | 0.95 | 0.33 | -0.14 | 0.98 |
| year2018 P. mesoamericanus / year2019 S. parvidens | 0.93 | 0.29 | -0.22 | 0.96 |
| year2019 P. mesoamericanus / year2017 S. parvidens | 2.39 | 1.21 | 1.71 | 0.24 |
| year2019 P. mesoamericanus / year2018 S. parvidens | 1.02 | 0.26 | 0.09 | 0.98 |
| year2019 P. mesoamericanus / year2019 S. parvidens | 1 | 0.21 | 0 | 1 |
| year2017 S. parvidens / year2018 S. parvidens | 0.43 | 0.23 | -1.6 | 0.24 |
| year2017 S. parvidens / year2019 S. parvidens | 0.42 | 0.21 | -1.71 | 0.24 |
| year2018 S. parvidens / year2019 S. parvidens | 0.98 | 0.25 | -0.08 | 0.98 |

*Additional Methods and Results for Hemoplasma Diagnostics*

PCR-positive amplicons for the 16S rRNA gene of hemoplasmas were purified using the QIAquick PCR Purification Kit (Qiagen). Positive amplicons were directly sequenced in both directions using the primers used for PCR and internal primers at Psomagen followed by NCBI BLASTn. Phylogenetic analyses and genotype assignments of hemoplasmas followed previously established methods (Volokhov *et al*. 2017; Becker *et al.* 2020a).

For *Desmodus rotundus*, positives belonged to the previously established genotypes VBG1, VBG2, and VBG3 (Volokhov *et al*. 2017), all of which are largely specific to *D. rotundus* with rare infection in *Pteronotus* species (Becker *et al.* 2020a). For *Sturnira parvidens*, we primarily detected hemoplasmas belonging to the previously identified SP1 genotype (groups A–C), largely specific to *S. parvidens* but also detected occasionally in another fruit bat, *Artibeus lituratus* (Becker *et al.* 2020a). However, one individual of *S. parvidens* had a novel genotype (SP2; GenBank accession number OQ308927) showing 97% similarity to the APH3 genotype (Becker *et al.* 2020a), previously observed only in the fruit bat *Artibeus intermedius* (Becker *et al.* 2020a). Lastly, for *Pteronotus mesoamericanus*, we observed infection most commonly with the PPM1 genotype (previously found in this same species; Becker *et al.* 2020a), although we also detected an infection with the SP1 genotype (more commonly found in *S. parvidens*) as well as a novel genotype (PPM2; GenBank accession number OQ308895) with 97% similarity to VBG1 (Becker *et al.* 2020a), previously only found in *D. rotundus* and the closely related *Pteronotus fulvus* (Becker *et al.* 2020a). *P. mesoamericanus* was also infected by a novel non-hemotropic *Mycoplasma*, the *M. moatsii*–like genotype 4 (GenBank accession number OQ308889); other *M. moatsii*–like genotypes have been previously identified in *D. rotundus*, *P. mesoamericanus*, and the insectivorous bats *Myotis pilosatibialis* and *Rhynchonycteris naso* from Belize (Becker *et al.* 2020a; Volokhov *et al*. 2017).

*Works Cited*

Becker, D. J., Speer, K. A., Brown, A. M., Fenton, M. B., Washburne, A. D., Altizer, S., Streicker, D. G.,

Plowright, R. K., Chizhikov, V. E., Simmons, N. B., & Volokhov, D. V. (2020a). Ecological and

evolutionary drivers of haemoplasma infection and bacterial genotype sharing in a Neotropical bat community. *Molecular Ecology*, 29(8), 1534–1549.

Volokhov, D. V., Becker, D. J., Bergner, L. M., Camus, M. S., Orton, R. J., Chizhikov, V. E., Altizer, S.

M., & Streicker, D. G. (2017). Novel hemotropic mycoplasmas are widespread and genetically diverse in vampire bats. *Epidemiology and Infection*, *145*(15), 3154–3167.
